# Metabolic Trans-Omic Analysis Reveals Key Regulatory Disruption of Energy Metabolism in Alzheimer’s Disease

**DOI:** 10.1101/2025.09.26.678758

**Authors:** Tomoharu Katayama, Hikaru Sugimoto, Keigo Morita, Hirohisa Watanabe, Shinya Kuroda

## Abstract

Alzheimer’s disease (AD), a leading cause of dementia, has been recognized as a disease with profound metabolic dysregulation. However, a systems-level view of metabolic regulation across multiple omic modalities in AD remains elusive. Here, we integrated public multi-omic datasets (transcriptome, proteome, and metabolome) from the dorsolateral prefrontal cortex of AD patients and controls. By leveraging existing molecular biological knowledge, we reconstructed a multi-layered metabolic regulatory network to systematically map the potential interplay among mRNAs, proteins, and metabolites in AD. Our analysis revealed a putative coordinated downregulation of energy producing pathways, including the TCA cycle, oxidative phosphorylation, and ketone body metabolism, driven by reduced enzyme abundance and inhibitory allosteric effects. In contrast, the glycolysis pathway appeared to be influenced by opposing enzymatic and allosteric regulations. These findings highlight key metabolic dysregulations that may contribute to the bioenergetic deficits in AD.

## Introduction

Alzheimer’s disease (AD) is a leading cause of dementia and affects tens of millions of people worldwide currently (Breijyeh & Karaman, 2020). It is pathologically characterized by senile plaques, neurofibrillary tangles, and cerebral atrophy (Breijyeh & Karaman, 2020). Many risk factors are involved in the development of this disease: aging, genetic factors, vascular disease, infections, environmental factors, and metabolic disorders (Scheltens et al., 2021). Among these hallmarks, some studies have focused on the role of altered metabolism in AD. One well-known abnormality that appears early in the progression of dementia is downregulation of glucose metabolism (Yan et al., 2020). In the clinical settings, fluorodeoxyglucose (FDG)-positron emission tomography (PET) is used to detect the changes in glucose metabolism in brain and to distinguish between different types of dementia (de la Monte & Tong, 2014). FDG-PET detects reduced glucose consumption in the parietal-temporal area, medial temporal lobes, and posterior cingulate cortices during early AD, which later extends to the frontal lobes and subcortical areas (Mosconi et al., 2009; Szablewski, 2021). One of the causes for the dysregulation of glucose metabolism is the downregulation of GLUT1 and GLUT3, glucose transporters involved in glucose uptake into brain tissue. These transporters are downregulated in AD patients (Kyrtata et al., 2021; Liu et al., 2008; Simpson et al., 1994), correlating with the extent of amyloid beta (Aβ) accumulation and neurofibrillary tangles (An et al., 2018; Liu et al., 2008). Mitochondrial dysfunction is another key feature of AD pathology (Swerdlow, 2020). Studies of postmortem brain tissue have revealed impairments in the pyruvate dehydrogenase complex (PDHC) (Bubber et al., 2005; Parker et al., 1994) and mitochondrial respiratory chain complexes (Parker et al., 1994) in patients with AD compared to cognitively normal subjects. More recently, there has been growing interest in the link between type 2 diabetes mellitus and AD, because brain insulin resistance has been observed in AD brains. This pathology, defined as the failure of brain cells to respond to insulin, leads to synaptic dysfunction and impaired immune response (Arnold et al., 2018; Steen et al., 2005). Together, these metabolic abnormalities result in bioenergetic deficits, oxidative stress, chronic inflammation, and protein quality control failures in AD (Kumar et al., 2022).

Large-scale and multi-omic analyses have identified many genes, proteins, and metabolic pathways that are associated with AD (Beckmann et al., 2020; Leventhal et al., 2024; M. Wang et al., 2021). For example, co-expression network analysis of brain proteomic data has identified protein network modules associated with microglia, astrocytes, and sugar metabolism as the most strongly altered in AD (Johnson et al., 2020). Integrative multi-omic network analysis has shown that short-chain acylcarnitines/amino acids and medium/long-chain acylcarnitines are most associated with AD clinical outcomes, implicating regulatory roles of *ABCA1* and *CPT1A* (Horgusluoglu et al., 2022). More recently, machine learning techniques have been applied to the integration of multi-omic data and the identification of functional modules and key molecules related to AD (Moon & Lee, 2022; J. Wang et al., 2024). However, these studies predominantly focus on correlations-based analyses and biomarker discovery rather than elucidating the detailed mechanistic regulation by the physicochemical interactions among molecules across different modalities.

“Trans-omic analysis” can be a powerful approach for reconstructing a cross-hierarchical molecular regulatory network by integrating comprehensive quantitative data of molecules measured at multiple levels (gene expression level, protein abundance, and metabolite abundance) with existing molecular biological knowledge. Such an approach has been applied to study obesity in mouse models and identified regulatory changes in metabolic reactions in the liver and muscle in either the fasting state or after glucose administration in wild-type mice and *ob/ob* mice (Egami et al., 2021; Kokaji et al., 2022; Sugimoto et al., 2024).

Here, we reconstructed a trans-omic network to uncover regulatory changes in metabolic reactions between cognitively normal control subjects and AD patients. We used public multi-omic datasets (transcriptomic, proteomic, and metabolomic data) of the dorsolateral prefrontal cortex (DLPFC) from the Religious Order Study and the Memory and Aging Project (ROSMAP) study (Bennett et al., 2018, 2012). The metabolic trans-omic analysis revealed putative AD-associated dysfunction due to changes in enzyme abundance and allosteric regulation by metabolites in pathways related to energy production, including the glycolysis pathway, the TCA cycle, the oxidative phosphorylation pathway, and ketone body metabolism. Our study provides a systems-level insight into intracellular metabolic dysregulation that may underlie the pathology of AD.

## Results

### Overview of trans-omic analysis for metabolic reactions of Alzheimer’s disease

To understand metabolic dysfunction in AD, we constructed a trans-omic network that integrated transcriptomic, proteomic, and metabolomic data from postmortem human brain tissue. We primarily used public multi-omic datasets (Batra et al., 2023; De Jager et al., 2018; Johnson et al., 2020) obtained from the DLPFC collected from cognitively normal individuals and patients with AD in the ROSMAP study (Bennett et al., 2018, 2012). Although pathological changes in AD are prominent in other brain areas, including temporal and parietal lobes, our study focused on the DLPFC due to the unique availability of comprehensive and matched multi-omic data for this region. The transcriptome was measured by bulk RNA sequencing (NGS), the proteome by tandem mass tag (TMT)-MS, and the metabolome by UPLC-MS/MS (Fig. 1).

**Figure 1.**
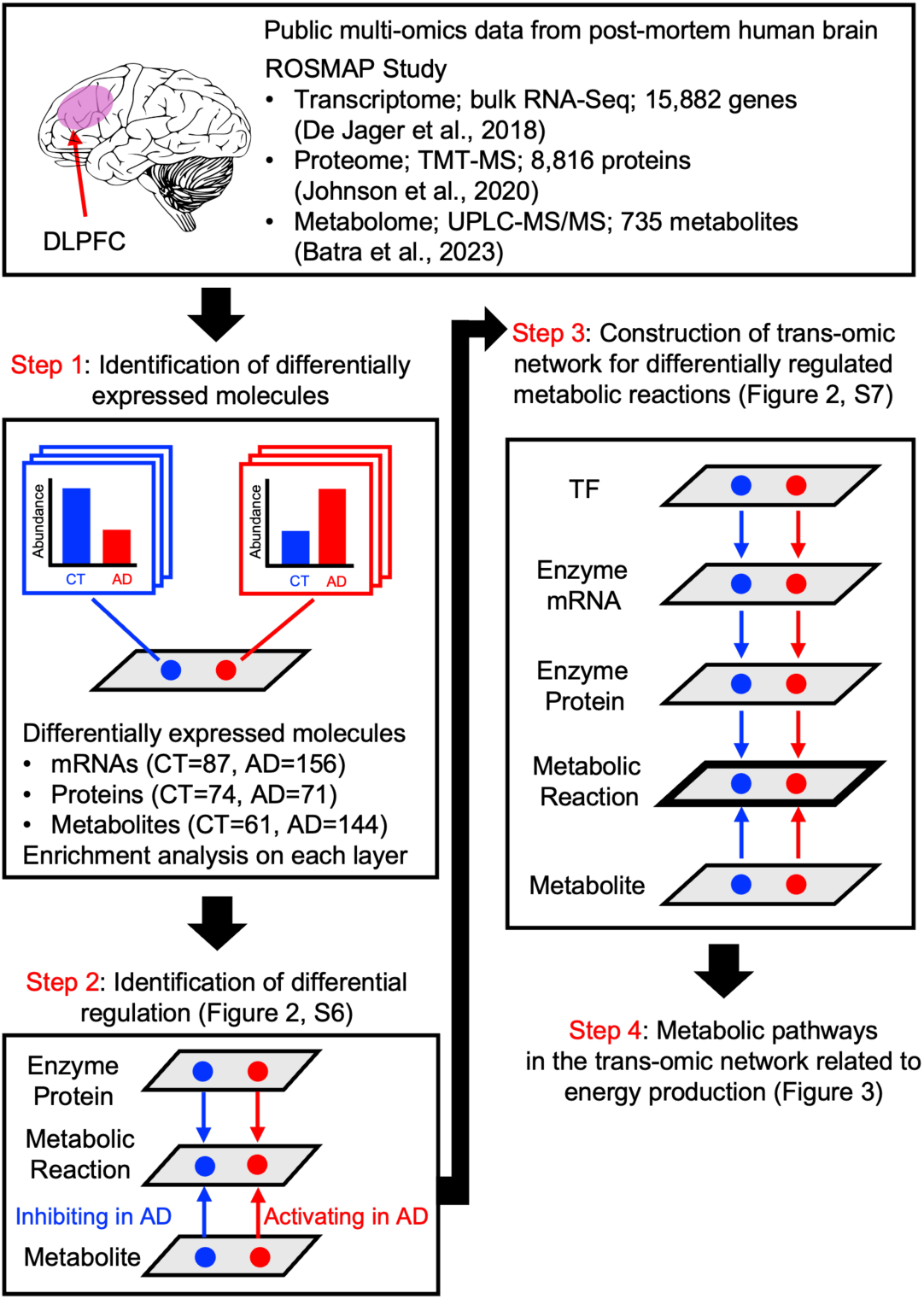
Overview of trans-omic analysis for metabolic reactions in Alzheimer’s disease. The illustration shows how public multi-omic data from the DLPFC of cognitively normal subjects (CT) and patients with Alzheimer’s disease (AD) from the ROSMAP study were used to generate a metabolic trans-omic network (Steps 1-3) and to perform detailed analysis of energy production pathways (Step 4). Red indicates differentially expressed molecules with increased expression or abundance, regulatory events that activate metabolic reactions, or increased activity of a metabolic reaction. Blue indicates differentially expressed molecules with decreased expression or abundance, regulatory events that inhibit metabolic reactions, or reduced activity of a metabolic reaction. The sample size for each analysis: n = 87 (CT group) and n = 156 (AD group) for transcriptomics, n = 74 (CT group) and n = 71 (AD group) for proteomics, and n = 61 (CT group) and n = 144 (AD group) for metabolomics.

We identified differentially expressed genes (DEGs), proteins (DEPs), and metabolites (DEMs) in patients with AD compared to cognitively normal controls using each omic dataset (Fig. 1, Step 1). To focus on metabolic regulation, we used KEGG (Kanehisa et al., 2025) and BRENDA (Chang et al., 2021) to match enzymes and metabolites to metabolic reactions and then determined AD-associated differences in metabolic regulation (“differential regulation”) (Fig. 1, Step 2). We also identified the enzyme mRNAs and transcription factors (TFs) that may be involved in the metabolic changes in AD (Zou et al., 2024). We then reconstructed a metabolic trans-omic network through integrating all these molecular relationships spanning multi-omic modalities (Fig. 1, Step 3). We further analyzed key metabolic pathways in the trans-omic network related to energy production, including glycolysis, the TCA cycle, oxidative phosphorylation, and ketone body metabolism (Fig. 1, Step 4).

### Identification of differentially expressed molecules of AD

We accessed the omic datasets and associated clinical, demographic, and postmortem neuropathological data for each individual from the ROSMAP study through the AD Knowledge Portal (https://adknowledgeportal.synapse.org/). A detailed summary of the samples used in this study, including the overlap across the three omicmodalities, is provided in Fig. S1. To ensure the representativeness of this cohort, we compared the ROSMAP study dataset with other public datasets (MSBB study and Wan et al., 2020 for transcriptomic validation; AMP-AD Diverse Cohorts for proteomic validation; Batra et al., 2024 and Novotny et al., 2023 for metabolomic validation), confirming significant correlations or overlap in AD-related changes across independent datasets (Fig. S2). Furthermore, to address potential alterations in the cell type proportions associated with AD, we identified overlapping individuals between our bulk omic datasets and a previously published single-nucleus transcriptomic dataset (Green et al., 2024). For the overlapping samples, we calculated major cell type proportions and statistically compared them between cognitively normal subjects and patients with AD (Fig. S3). In the transcriptomic data, we have confirmed significantly increased proportion of oligodendrocytes and decreased proportion of inhibitory neurons in patients with AD. In the metabolomic data, we found significantly decreased proportion of inhibitory neurons in patients with AD.

We defined the cognitively normal control (CT) group and the AD group based on clinical diagnosis and neuropathological diagnoses (Braak Stage and CERAD score) (see also Fig. S1A and Methods section) (Bennett et al., 2018, 2012). We defined 946 significantly increased mRNAs and 917 decreased mRNAs in AD using the results of the differential expression analysis from the RNA-seq Harmonization Study (Wan et al., 2020) (Synapse id: syn8456721). We identified 657 significantly increased proteins and 854 decreased proteins in AD using the proteomic data in the ROSMAP Study (id: syn21266454) (Johnson et al., 2020). We identified 79 significantly increased metabolites and 78 decreased metabolites in AD using the metabolomic data from the ROSMAP study (id: syn25985690) (Batra et al., 2023).

Additionally, we conducted enrichment analyses for each omic dataset (Fig. S4D), highlighting upregulation of nitrogen metabolism and downregulation of ribosomal function. Furthermore, we constructed protein-protein interaction networks for the increased and decreased DEPs using the STRING database (Szklarczyk et al., 2023) (Fig. S5 and Supplementary Data 2). The result indicated downregulated translation, both in the nucleus and in the mitochondria. We also visualized the top 20 DEGs, DEPs, and DEMs with log_2_FC values (Fig. S6 and Supplementary text). Of note, the abundance of ATP was not quantified in the metabolomic data from the ROSMAP study (Batra et al., 2023).

### Relationship between proteomic changes and gene expression changes in AD

To examine the relationship between DEGs and DEPs, we investigated the expression changes of the genes encoding each DEP (Fig. S7A). Among the increased DEPs, 76 DEPs were encoded by significantly upregulated genes, and 15 DEPs were encoded by significantly downregulated genes. Among the decreased DEPs, 70 DEPs were encoded by genes with significantly decreased expression, and 12 DEPs were encoded by genes with significantly increased expression. Thus, many of the DEPs did not have coordinated expression changes with their encoding genes (Fig. S7B). This result is consistent with previous studies using postmortem human transcriptomic and proteomic data (Johnson et al., 2022; Tasaki et al., 2022).

### Identification of differentially expressed transcription factors

We identified TFs potentially involved in regulating the expression changes of the DEGs using ChIP-Atlas, a database that integrates and provides ChIP-seq data (Zou et al., 2024). No TFs were predicted to regulate genes with significantly decreased expression; 19 TFs were identified as regulators of genes with significantly increased expression (Table S1). To determine the TFs with differential activity in AD, we investigated whether the predicted TFs were encoded by DEGs or were DEPs (Fig. S7C). We identified five TFs that were significantly increased at either the transcriptomic level (CTCF, MAX, and ZNF143) or the proteomic level (BRD4 and RAD21). We defined these 5 TFs as “differentially expressed transcription factors” (DETFs). Recent findings have shown the protective role of BRD4, which inhibits transcriptional expression of AD-risk genes (Matuszewska et al., 2022; Nikkar et al., 2022; Sun et al., 2024).

### Identification of metabolic enzymes among the differentially expressed proteins

To focus on the proteomic changes in AD that are relevant to cellular metabolism, we identified metabolic enzymes among the DEPs according to their designations as metabolic enzymes in the “Metabolic pathways” of the KEGG PATHWAY Database (Kanehisa et al., 2025). Among the 657 significantly increased proteins, 59 are metabolic enzymes. Among the 854 significantly decreased proteins, 70 are metabolic enzymes (Table S2).

### Identification of metabolites that are substrates, products, or allosteric regulators of metabolic reactions

Metabolic reactions are regulated not only by the abundance of the metabolic enzymes that catalyze the reactions, but also by the abundance of substrates and products, as well as by allosteric regulators. Using the BRENDA database (Chang et al., 2021), we identified allosteric regulators among the DEMs: 46 were increased in AD and 26 were decreased in AD (Fig. 2A). These 72 allosteric regulatory DEMs affected 318 enzymes. Using the KEGG REACTION database (Kanehisa et al., 2025), we also identified substrates and products of metabolic reactions among the DEMs: 50 substrates or products were increased in AD and 33 were decreased in AD, affecting 727 enzymes (Fig. 2B).

**Figure 2.**
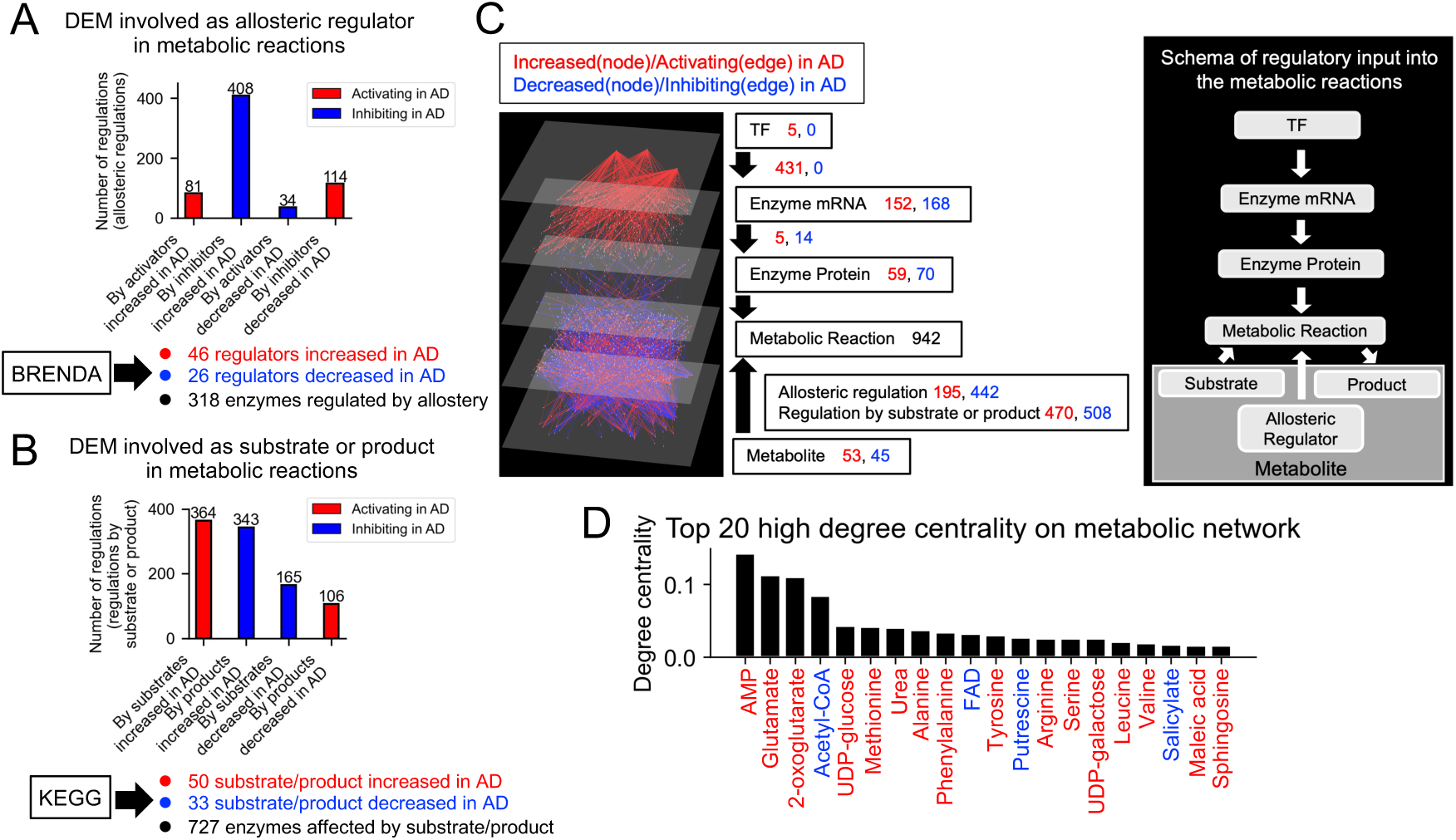
Construction of a trans-omic network for differentially regulated metabolic reactions. (A) The number of allosteric regulators among DEMs and the number of allosteric regulations mediated by them on metabolic enzymes identified with BRENDA. Some reactions are regulated by more than one DEM and some DEMs regulate multiple enzymes, thus the number of regulations exceeds the number of enzymes identified with BRENDA and the number of DEMs. (B) The number of substrates or products among DEMs and the number of regulations by them on metabolic enzymes identified with KEGG. Some reactions involve multiple substrates or yield multiple products and some substrates and products occur in multiple reactions, thus the number of regulations exceeds the number of enzymes identified with KEGG and the number of DEMs. (C) The trans-omic network for differentially regulated metabolic reactions (left). Nodes and edges indicate differentially expressed molecules and differential regulations involved in metabolic reactions, respectively. Schema of regulatory input into the metabolic reactions (right). TFs control metabolic enzyme mRNA abundance through gene regulation. Metabolite acts as substrates, products, and allosteric regulators in metabolic reactions. See Fig. S8 and Methods section for additional details. (D) Top 20 molecules with high degree centrality on the metabolic network. The metabolic network consisted of the nodes and edges from the “Enzyme Protein”, “Metabolic Reaction”, and “Metabolite” layers of the trans-omic network. Those shown in red text were increased in AD; those in blue were decreased in AD.

### Construction of a trans-omic network for differentially regulated metabolic reactions

We constructed a metabolic trans-omic network for differentially regulated metabolic reactions in AD (Fig. 2C and Supplementary Data 1). This network integrates five interconnected components organized into corresponding layers: (1) DETFs in the “TF layer”, (2) DEGs encoding metabolic enzymes in the “Enzyme mRNA layer”, (3) DEPs functioning as metabolic enzymes in the “Enzyme Protein layer”, (4) metabolic reactions differentially regulated by enzymes or metabolites in the “Metabolic Reaction layer”, and (5) DEMs involved in the metabolic reactions as allosteric regulators, products, or substrates in the “’Metabolite layer”. The adjacent layers are connected by edges based on the regulatory relationships between the molecules. The right panel of Fig. 2C shows the schema for the construction of the trans-omic network, corresponding to the five layers in the left panel. Specifically, the interlayer connections from the “Metabolite layer” to the “Metabolic Reaction layer” consisted of regulatory connections mediated by allosteric regulators, substrates, or products of each metabolic reaction (Egami et al., 2021; Kokaji et al., 2022). Activating regulation of metabolic reactions is shown as red edges, and inhibiting regulation is shown as blue edges (see also Fig. S8, and Methods section).

We analyzed the degree centrality of nodes in the metabolic reaction network that consisted of “Enzyme Protein layer”, “Metabolic Reaction layer”, and “Metabolite layer.” Degree centrality is a measure of the importance of a node in network theory (Koschützki & Schreiber, 2008). The degree centrality of a node is calculated by normalizing the number of edges connected to that node, with a higher value indicating greater influence within the network. AMP, which increased in the AD group, had the highest degree centrality (Fig. 2D). Glutamate, 2-oxoglutarate (α-ketoglutarate), and urea, which were increased in AD, were second, third, and seventh in degree centrality, respectively. Acetyl-CoA, which decreased in the AD group, ranked fourth.

### The metabolic dysregulations of energy metabolic pathways of AD

We extracted key metabolic pathways involved in energy production in the trans-omic network, the glycolysis pathway, the TCA cycle, oxidative phosphorylation, and ketone body metabolism. We examined detailed metabolic dysregulation in each pathway (Fig. 3).

**Figure 3.**
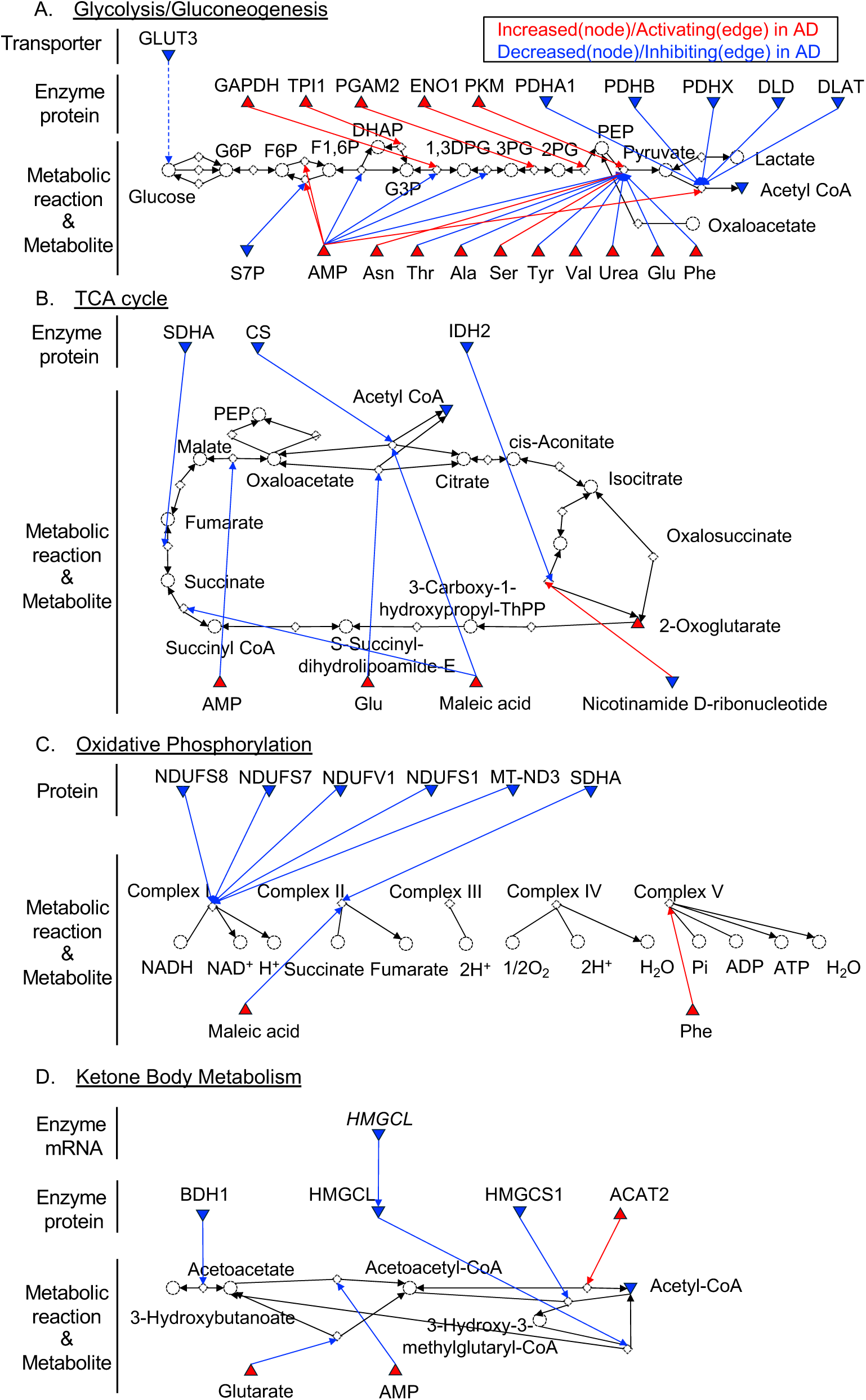
The metabolic dysregulations of energy metabolic pathways of AD. The trans-omic changes in the regulations of glycolysis (A), TCA cycle (B), oxidative phosphorylation (C), and ketone body metabolism (D). The information for each pathway were obtained from “glycolysis/gluconeogenesis” (hsa00010), “TCA cycle” (hsa00020), “oxidative phosphorylation” (hsa00190), and “butanoate metabolism” (hsa00650) in the KEGG database. Red upward triangles, increased molecules in AD; blue downward triangles, decreased molecules in AD. Red arrow, regulations that activate metabolic reactions; blue arrow, regulations that inhibit reactions. Unfilled circles with dashed outlines represent molecules that were not differentially expressed or were not measured. Unfilled diamond nodes with dashed outlines represent metabolic reactions. See Fig. S8 and Methods section for additional details.

In the glycolysis pathway, increased expression of TPI1, GAPDH, PGAM2, ENO1, and PKM would increase flux through the glycolysis pathway, whereas the increased metabolites S7P, AMP, threonine, alanine, tyrosine, valine, urea, glutamate, and phenylalanine would allosterically inhibit flux through the pathway. In addition, the expression of GLUT3, a transporter responsible for the basal uptake of glucose in neurons (Koepsell, 2020), is significantly decreased (Fig. 3A), as demonstrated in several previous studies (An et al., 2018; Liu et al., 2008; Simpson et al., 1994). However, the mRNA expression of *GLUT3* did not show a significant change in AD in the transcriptomic dataset we used.

Previous studies have reported downregulation of PDHC in AD patients (Jankowska-Kulawy et al., 2022; Jia et al., 2023). Consistently, we found that five of the six proteins comprising the complex, PDHA1, PDHB, PDHX, DLD and DLAT, were significantly decreased. In addition, acetyl-CoA, was significantly decreased in AD. The previous studies (Jankowska-Kulawy et al., 2022; Jia et al., 2023) and our findings suggest that reduced PDHC activity may decrease the production of acetyl-CoA in AD.

In the TCA cycle, most regulations were inhibitory (Fig. 3B). Three metabolic enzymes were significantly decreased: SDHA, CS, and IDH2. Increases in AMP, glutamate, and maleic acid resulted in allosteric inhibition of metabolic reactions in the TCA cycle. Only a reduction in nicotinamide D-ribonucleotide exhibited an allosteric activating effect. Among the five measured metabolites (citrate, aconitate, 2-oxoglutarate, fumarate, and malate) that are substrates and products in the TCA cycle, 2-oxoglutarate (α-ketoglutarate) showed a significant increase in abundance.

In oxidative phosphorylation, most of the regulatory changes in AD resulted in inhibition of metabolic reactions (Fig. 3C). Five subunits of complex I in the respiratory chain were decreased: NDUFS8, NDUFS7, NDUFV1, NDUFS1, and MT-ND3. This result is consistent with other postmortem studies of human brains which have shown reduced expression of genes encoding subunits of the electron transport chain in posterior cingulate cortex, middle temporal gyrus, and the hippocampus of AD patients (Liang et al., 2008). The activity of complex II was also negatively impacted by the decrease in SDHA and an increase in the allosteric inhibitor, maleic acid. The increase in phenylalanine, an allosteric regulator of complex V, provided the only activating regulation.

Ketone bodies are the second-most utilized energy source in the brain after glucose (Jensen et al., 2020). Ketone bodies are synthesized in the liver in the form of 3-hydroxybutyrate and acetoacetate, which are transported to the brain through the circulation (Jensen et al., 2020). Ketone bodies undergo several metabolic reactions and are converted to acetyl-CoA as an energy source in mitochondria (Jensen et al., 2020). Most of the AD-associated changes resulted in inhibition of ketone body metabolism (Fig. 3D). Among the metabolic enzymes involved in ketone body metabolism, BDH1, HMGCL, and HMGCS1 were downregulated; only ACAT2 was upregulated. The increase in glutarate and AMP resulted in inhibitory allosteric regulation. Collectively, these results suggested that reduced ketone body metabolism contributed to the decrease in acetyl-CoA abundance in AD.

Through this examination of the changes in the regulation of core metabolic pathways involved in energy production, we found putative antagonistic regulation by enzymes and allosteric regulators in the glycolysis pathway. On the other hand, the TCA cycle, oxidative phosphorylation, and ketone body metabolism can be downregulated in AD due to the reduced enzymes and inhibitory allosteric effects. Here, we demonstrate the systems-level view of the potential dysregulated energy production of AD in terms of the metabolic trans-omic network.

## Discussion

In this study, we identified differentially expressed molecules from public datasets of transcriptome, proteome, and metabolome obtained from the ROSMAP study and used the results to construct a metabolic trans-omic network in AD. We then performed a detailed analysis of the AD-associated changes in pathways related to energy production. Above all, we summarized the metabolic dysregulations observed in the brain of AD (Fig. 4).

**Figure 4.**
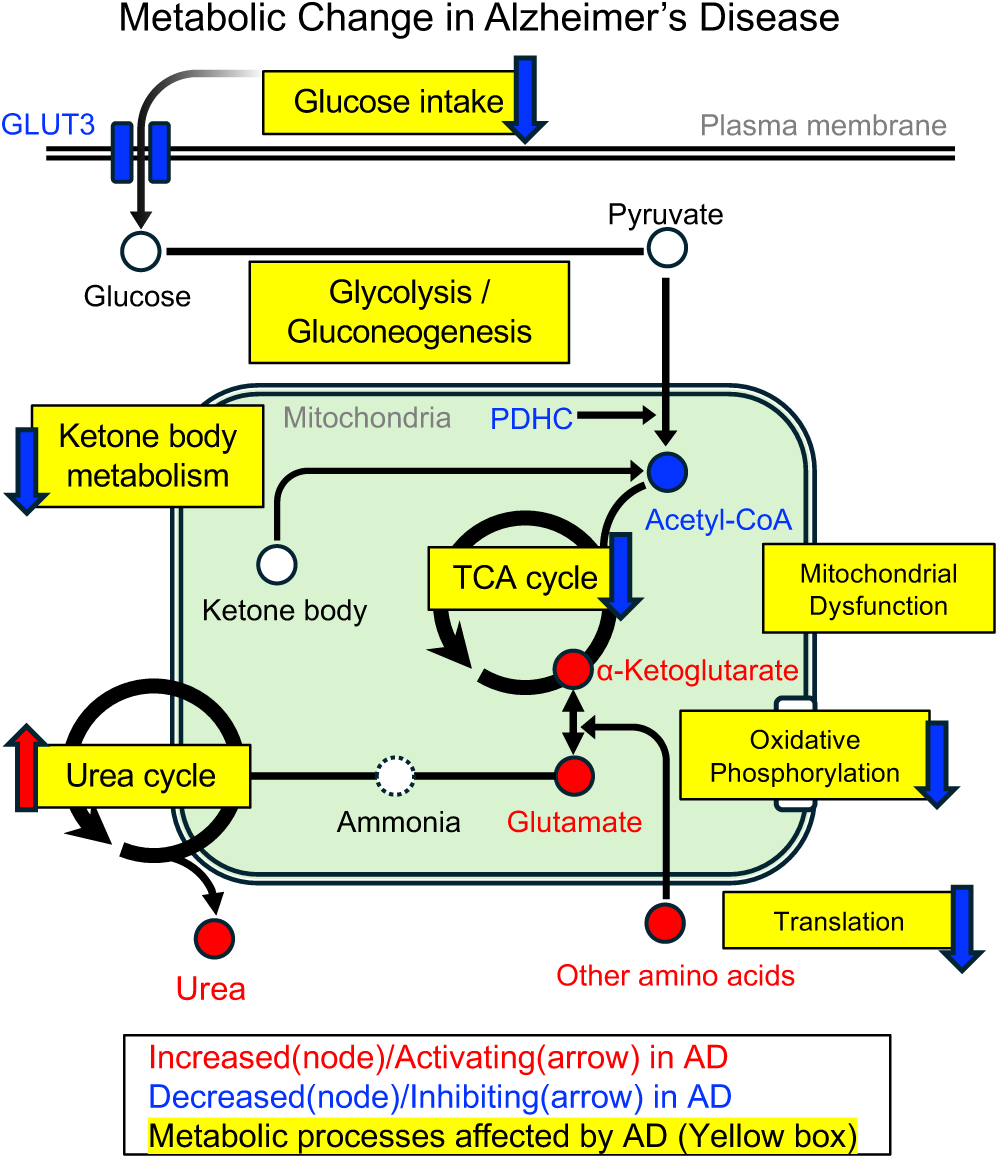
Models of potentially dysregulated metabolic processes in AD. Diagrams of dysregulated metabolic pathways associated with the progression of AD. Metabolic processes affected by AD are highlighted in yellow boxes. Red, increased (circles) or activating (arrows) in AD; blue, decreased (circles and GLUT3) or inhibiting (arrows) in AD.

Several key factors that underlie the mechanism of dysregulated energy metabolism have been identified in many previous studies (An et al., 2018; Arnold et al., 2018; Bubber et al., 2005; Kyrtata et al., 2021; Liu et al., 2008; Parker et al., 1994; Simpson et al., 1994; Steen et al., 2005; Swerdlow, 2020; Yan et al., 2020), including downregulated glucose uptake and utilization, impaired mitochondrial functions, dysfunctional enzyme proteins, and brain insulin resistance, among others. However, details of the regulatory relationships between the molecules belonging to different modalities have been poorly understood. Here, we demonstrated a potential mechanistic picture of dysregulated core energy metabolism in AD (Fig. 4): (1) inhibition of the TCA cycle, oxidative phosphorylation, and ketone body metabolism by allosteric inhibition and downregulated enzyme proteins, (2) antagonistic regulation by enzymes and allosteric regulators in the glycolysis pathway.

As previously reported (Yu et al., 2022), mitochondrial dysfunction accompanies the progression of AD, which may lead to downregulation of TCA cycle, oxidative phosphorylation, and ketone body utilization pathways, as we confirmed in this study, since these metabolic reactions occur within mitochondria. Additionally, decreased expression of the subunit proteins that make up PDHC and allosteric inhibition to the glycolysis pathway may further reduce acetyl-CoA levels, resulting in a greater decrease in the ability to produce enough ATP. This energetic shortage appears to be partially compensated by upregulation of glycolytic enzymes, but this compensatory mechanism could be inhibited by the reduced glucose uptake through GLUT3 in neurons. This hypothesis is supported by the previous study, which has suggested an upregulated glycolysis pathway in astrocytes as a compensatory mechanism to provide the energy source for impaired neurons during AD progression (Fu et al., 2015).

The DEM analysis revealed significant increases in 12 amino acids (Glu, Tyr, Thr, Phe, Asn, Ser, Arg, Val, Leu, Ala, Met, and Lys), 2-oxoglutarate, and urea. In addition, we identified ASL, an enzyme involved in the urea cycle (Hansmannel et al., 2010), as significantly increased in AD. Thus, our results highlighted the potential involvement of nitrogen metabolism dysregulation in AD and suggested that the observed accumulation of urea may result from enhanced flux through the urea cycle within the DLPFC. This could reflect a response to elevated amino acid levels, potentially linked to compromised translation and protein turnover. This hypothesis is also supported by the results of our PPI network analysis and the low correlation between mRNA expression and protein expression.

Previous studies have demonstrated that transcripts of ribosomal proteins are significantly decreased in the dentate gyrus in AD patients (Hernández-Ortega et al., 2016). In another recent research, changes in the proteome are important for understanding AD pathology, because approximately half of the network modules identified in protein networks were not found in RNA networks (Johnson et al., 2022). About nitrogen metabolism in AD, a previous study has demonstrated that astrocytic urea metabolism induced in AD has a beneficial effect on Aβ detoxification. However, this metabolic process has been found to induce memory impairment through producing toxic byproducts, ammonia and H_2_O_2_, and GABA (Ju et al., 2022).

### Limitations of this study

Several methodological limitations of our study should be acknowledged. Our analysis did not distinguish between cell types. Brain tissue contains various types of cells: neurons, astrocytes, oligodendrocytes, and microglia. The genes and proteins expressed in each cell type differ, as do the metabolic reactions that occur within each cell (Bonvento & Bolaños, 2021; Magistretti & Allaman, 2015). We primarily observed a decreased proportion of inhibitory neurons in the transcriptomic and metabolomic datasets (Fig. S3A and S3C). Additionally, previous studies has reported that Braak stage negatively correlated with the proportion of oligodendrocytes and positively correlated with the proportions of microglia and astrocytes (Hannon et al., 2024; Shireby et al., 2022). Therefore, our results could partly reflect changes in tissue composition associated with the progression of AD. A recent study using the snRNA-seq technique has demonstrated that in excitatory neurons downregulated DEGs that were correlated with NFT burden were enriched for TCA cycle (Mathys et al., 2024). In addition, the glycolysis and oxidative phosphorylation modules were upregulated in astrocytes and microglia in AD group (Mathys et al., 2024).

All omics data used in this study were measured from postmortem brain tissues. The cause of death and events that occur immediately before death (such as resuscitation) can influence molecular profiles of postmortem tissue. Thus, we must consider potential biases related to these factors. In addition, post-mortem interval (PMI) can affect the results of the quantifications of molecules. Regarding the metabolomic dataset that we used, about 300 metabolites were found to be associated with PMI in the original research (Batra et al., 2023). Thus, we included PMI as a covariate in the differentially expressed analysis.

In this study, we analyzed the DLPFC because that was the region with sufficient datasets. However, initial pathological changes of AD typically appear in the medial temporal lobe. Therefore, trans-omic analyses should also be conducted in this region. It is also possible that the results of this study reflect changes that compensate for metabolic alterations arising in other regions. However, at present, the number of large-scale proteomic and metabolomic datasets measured from human postmortem brains is limited.

Interpretation of regulatory relationships among altered molecules within the trans-omic network remains challenging because they are often highly complex and densely interconnected. For example, lipid metabolism is widely recognized as playing a critical role in AD pathology (Baloni et al., 2020; Horgusluoglu et al., 2022; Varma et al., 2021). However, our analytical framework did not allow for a sufficiently detailed dissection of lipid metabolic dysregulation. In addition, the trans-omic network analysis is sensitive to predefined analytical settings, including statistical thresholds for defining differentially expressed molecules and sample size. Therefore, our findings should be interpreted as an observational systems-level insights rather than definitive mechanistic conclusions into AD pathology. Further verification using independent and larger cohorts, as well as experimental validation will be essential in future studies.

## Materials & methods

### ROSMAP study

Public multi-omic datasets measured from postmortem human brain samples from the Religious Orders Study and Memory and Aging Project (ROSMAP) study were primarily used in this study. (https://adknowledgeportal.synapse.org/Explore/Studies/DetailsPage/StudyDetails?Study=syn3219045#StudyDescription) (Bennett et al., 2018, 2012). All data were obtained through the AD Knowledge Portal (https://adknowledgeportal.synapse.org/). The Religious Orders Study (ROS) and the Memory and Aging Project (MAP) are long-term studies that focus on memory, motor function, and functional problems in older adults. ROS includes Catholic nuns, priests, and brothers aged 65 and older; MAP includes older adults around Chicago. Both studies include annual clinical assessments, self-reports, and postmortem brain donations. Diagnosis of dementia is made using the same procedure in both studies, and final clinical diagnosis and pathological assessment are made after death. Further details of the studies can be found in (Bennett et al., 2018, 2012).

### RNA sequencing

For transcriptomic data, the results of differential expression analysis from the RNAseq Harmonization Study (De Jager et al., 2018) were used. For detailed methods of RNA-seq and differential expression analysis, see (https://adknowledgeportal.synapse.org/Explore/Studies/DetailsPage/StudyDetails?Study=syn3219045#Methods-GeneExpressionRNAseq) and (https://adknowledgeportal.synapse.org/Explore/Studies/DetailsPage/StudyDetails?Study=syn9702085#StudyDescription).

### Proteomic analysis

For the proteomic data, proteomic analysis in the ROSMAP study was used (Johnson et al., 2020). In that study, brain tissue samples from the ROSMAP cohort (n = 400) were tagged using the TMT 10-plex kit. Further details of the TMT-MS experiment are available at (https://adknowledgeportal.synapse.org/Explore/Studies/DetailsPage/StudyDetails?Study=syn3219045#Methods-ProteomicsTMTquantitation). For information on batch effect correction, outlier and covariate adjustment of the raw data, refer to (Johnson et al., 2020).

### Metabolomic analysis

For metabolomic data, UPLC-MS/MS analysis in the ROSMAP Study was used, which were generated at Metabolon and preprocessed by the Alzheimer’s Disease Metabolomics Consortium (Batra et al., 2023). Further details of the UPLC-MS/MS experiments and data normalization can be found at (https://adknowledgeportal.synapse.org/Explore/Studies/DetailsPage/StudyDetails?Stud y=syn3219045#Methods-MetabolomicsMetabolonHD4).

### Validation analysis

To ascertain how the choice of the ROSMAP Study as the data source could influence our analyses, the ROSMAP Study data were compared with other datasets or findings from specific published studies (Fig. S2). For transcriptomic validation, RNA-seq data from the Mount Sinai Brain Bank (MSBB) study (https://www.synapse.org/Synapse:syn10526259) were used. Log_2_ fold-change values were calculated for each gene by taking the log_2_ ratio of the mean expression in the AD group to that in the CT group, and Pearson correlation coefficients were calculated between the two datasets (Fig. S2A). In addition, we compared our ROSMAP-derived DEGs with those identified in a large-scale meta-analysis of 2,114 samples across seven brain regions using a fixed-effects model (q-value < 0.05) (Wan et al., 2020) (Fig. S2B, C). For validation of proteomic data, TMT-MS results from the Accelerating Medicines Partnership Alzheimer’s Disease Diverse Cohorts Study (AMP-AD DiverseCohorts) were used (https://www.synapse.org/Synapse:syn55249985) (Reddy et al., 2024).

Correlation analysis was performed as described above (Fig. S2D). We additionally calculated a separate correlation coefficient for the subset of metabolic enzymes involved in the energy metabolism analysis shown in Fig. 3. To assess concordance in a threshold-independent manner, we performed rank-rank hypergeometric overlap (RRHO) analysis (Cahill et al., 2018; Plaisier et al., 2010) (Fig. S2E). For RRHO, proteins were ranked by -log_10_(q-value or p-value) multiplied by the sign of the log_2_ fold-change calculated in each study. For metabolomic validation, we compared our results with previous studies (Maffioli et al., 2022; Paglia et al., 2016; Xu et al., 2016) (Fig. S2F) and previous metabolomic datasets measured by UPLC-MS/MS platform (Batra et al., 2024; Novotny et al., 2023). RRHO analyses were performed to evaluate concordance in ranked metabolite changes as described above (Fig. S2G).

### Cell type proportions analysis

A public single-nucleus transcriptomic dataset (Green et al., 2024) (id: syn31512863), which was obtained from DLPFC tissue of 465 subjects in the ROSMAP cohort, was used to assess the changes in cell type proportions associated with AD. For each omic data used in our analyses, we identified overlapping subjects and calculated the following cell type proportions for the matched samples: astrocytes, excitatory neurons, inhibitory neurons, microglia, oligodendrocytes, and OPCs. We statistically compared cell type proportions between control subjects and patients with AD using a two-tailed t-test, with the significance threshold set at p-value < 8.3 × 10^-3^ (Bonferroni correction).

### Definition of CT group and AD group

The CT group consisted of subjects whose clinical diagnosis was categorized as “No cognitive impairment”, with a Braak Stage of III or lower and a CERAD score indicating “no AD” or “possible AD”. The AD group comprised subjects whose clinical diagnosis was categorized as “Alzheimer’s dementia and no other cause of cognitive impairment”, with a Braak Stage of IV or higher and a CERAD score indicating “probable AD” or “definite AD”.

### Identification of differentially expressed molecules between CT and AD

To define the differentially expressed genes (DEGs) in AD compared to CT, the results of the differential expression analysis from the RNA-seq Harmonization Study were used (https://www.synapse.org/Synapse:syn8456721). In the study, the adjusted p-value and log2FC value of each gene in the two groups were calculated using the count data from RNA sequencing of postmortem brains (DLPFC) from 634 subjects (156 AD and 87 CT) in the ROSMAP Study. The values were calculated after adjusting for covariates such as age, gender, post-mortem interval (PMI), and RNA integrity number (RIN) score using the voom-limma package in R. In that analysis, genes with an adjusted p-value < 0.05 were defined as DEGs.

To identify the DEPs in AD compared to CT, the log2-transformed and normalized relative quantitative data (https://www.synapse.org/Synapse:syn21266454) measured by isobaric tandem mass tagging peptide labeling in the ROSMAP study (71 AD and 74 CT) were used. Following the exclusion of proteins with more than 50% missing values, ANCOVA was performed with gender and PMI as covariates for the remaining 8,816 proteins. Subsequently, the q-value and log2FC were calculated using the Storey’s method. Proteins with a q-value < 0.05 were defined as DEPs.

To identify the DEMs in AD compared to CT, the normalized quantified data (https://www.synapse.org/Synapse:syn25985690) measured using UPLC-MS/MS in the ROSMAP Study (144 AD and 61 CT) were used. Following the exclusion of metabolites with more than 50% missing values, ANCOVA was performed with gender and PMI as covariates for the remaining 735 metabolites. Subsequently, the q-value and log2FC were calculated using the Storey’s method. Metabolites with q-values < 0.05 were defined as DEMs.

### Enrichment analysis

Enrichment analyses were conducted on DEGs and DEPs using Metascape (version 3.5; https://metascape.org/gp/index.html#/main/step1) (Zhou et al., 2019). Enrichment analysis was conducted on DEMs using MetaboAnalyst (version 6.0; https://www.metaboanalyst.ca/) (Pang et al., 2024). The background was defined as all measured proteins for proteomic analysis, and all measured metabolites for metabolomic analysis.

### Inference of transcription factors for differentially expressed genes in DLPFC of AD

To infer the TFs involved in regulating the DEGs, we used the peak call data (TSS±1k) from ChIP-Atlas (https://chip-atlas.org/) (Zou et al., 2024). The “neural” cell type was selected, and a threshold of 200 was applied for the statistical significance calculated by MACS2. The p-value was calculated using Fisher’s exact test, and the Benjamini-Hochberg (BH) method was employed for FDR correction (q-value < 0.05).

### Construction of protein-protein interaction networks

In constructing the protein-protein interaction network (Fig. S4A, S4B), we included protein pairs with a score of 400 or more in the STRING data (Szklarczyk et al., 2023) as interacting pairs. We constructed the protein-protein interaction network by connecting interacting DEPs with edges.

### Construction of trans-omic network for differentially regulated metabolic reactions

DEPs and transcripts were matched using the Ensembl protein ID (Martin et al., 2023). Among the DEPs, proteins that were categorized as metabolic enzymes in the “metabolic pathways” of the KEGG PATHWAY database (Kanehisa et al., 2025) were identified. Using BRENDA (Chang et al., 2021), the allosteric regulators belonging to DEMs were identified. Using KEGG REACTION data, the DEMs that are involved as substrates or products in metabolic reactions were identified. By integrating the above information, we identified DEPs that act as metabolic enzymes, the metabolic reactions in which they are involved, and the DEGs encoding these proteins, as well as DEMs and the metabolic reactions in which these metabolites are involved as allosteric regulators or as products or substrates. To identify DETFs in AD, we examined which the inferred TFs were encoded by DEGs or appeared as the DEPs. All these data were used to construct a 5-layered (DETFs, DEGs, DEPs, metabolic reactions, DEMs) trans-omic network for metabolic reactions in the DLPFC of AD patients.

The adjacent layers are connected by edges based on the regulatory relationships between the molecules on these layers. The “Enzyme mRNA” and “TF” layers were connected by red edges when mRNA expression increased and by blue edges when it decreased. The “Enzyme mRNA” and “Enzyme Protein” layers were connected by red edges when both gene and protein expression of enzymes increased, and by blue edges when gene and protein expression of enzymes decreased. The “Enzyme Protein” and “Metabolic Reaction” layers were connected by red edges when enzyme expression increased, and by blue edges when enzyme expression decreased. For the connection between “Metabolic Reaction” and “Metabolite” layer, if a metabolite shows an increase in expression and also has an allosteric effect of activating the metabolic reaction it controls, it is colored red; if it shows an increase in expression and also has an allosteric effect of inhibiting the metabolic reaction it controls, it is colored blue; if it shows a decrease in expression and also has an allosteric effect of activating the metabolic reaction it controls, it is colored blue; if it shows a decrease in expression and also has an allosteric effect of inhibiting the metabolic reaction it controls, it is colored red. If a metabolite shows an increase in expression and it is a substrate of the metabolic reaction, it is colored red; if it shows an increase in expression and it is a product of the metabolic reaction, it is colored blue; if it shows a decrease in expression and it is a substrate of the metabolic reaction, it is colored blue; if it shows a decrease in expression and it is a product of the metabolic reaction, it is colored red.

## Implementation

Statistical tests were performed using MATLAB 2023b (The Mathworks Inc; https://www.mathworks.com/products/matlab.html). Trans-omic network analysis and visualization were implemented in Python (version 3.12.2; https://www.python.org/), Pandas (version 1.5.3; https://pandas.pydata.org/), Networkx (version 3.1; https://networkx.org/), and Matplotlib (version 3.7.1; https://matplotlib.org/) packages.

## Resource availability Lead contact

Further information and requests for resources and reagents should be directed to and will be fulfilled by the Lead Contact, Shinya Kuroda.

## Materials availability

This study did not generate any new material.

## Data availability

All raw omics data used for this study are available through the AD Knowledge Portal (https://adknowledgeportal.org). The AD Knowledge Portal is a platform for accessing data, analyses, and tools generated by the Accelerating Medicines Partnership (AMP-AD) Target Discovery Program and other National Institute on Aging (NIA)-supported programs to enable open-science practices and accelerate translational learning. The data, analyses and tools are shared early in the research cycle without a publication embargo on secondary use. Data are available for general research use according to the following requirements for data access and data attribution (https://adknowledgeportal.synapse.org/Data%20Access).

## Code availability

We utilized and modified the algorithm that was developed in our previous study, iTraNet (Sugimoto et al., 2024). The code used for the analysis in this paper is available upon request.

## Acknowledgements

We thank Takashi Iwatsubo (The University of Tokyo), Kenichiro Sato (The University of Tokyo), Tadafumi Kato (Juntendo University), Katsuyuki Yugi (RIKEN), Junichi Maruyama (RIKEN), Kentaro Tomii (AIST), and Hafumi Nishi (Tohoku University), Tatsuhiko Tsunoda (The University of Tokyo) for valuable suggestions and discussions. We also thank our laboratory members for critically reading this manuscript.

This study was supported by the Japan Society for the Promotion of Science (JSPS) KAKENHI (JP21H04759), CREST, the Japan Science and Technology Agency (JST) (JPMJCR2123), JST ASPIRE (JPMJAP24B1), and by The Uehara Memorial Foundation. K.M. receives funding from a Grant-in-Aid for Early-Career Scientists (JP21K15342). H.W. was supported by AMED Grant Number JP21wm0425016.

The results published here are in whole or in part based on data obtained from the AD Knowledge Portal (https://adknowledgeportal.org/).

For the transcriptome data, we used the results of differential expression analysis based on RNA-seq measurements from ROSMAP participants as part of the RNA-seq Harmonization Study (https://www.synapse.org/Synapse:syn8456721). We utilized proteome data obtained from participants of the ROSMAP Study (https://www.synapse.org/Synapse:syn21266454). We used metabolome data acquired from ROSMAP Study participants (https://www.synapse.org/Synapse:syn25985690).

For validation of transcriptome data, we used the results of a differential expression analysis conducted using the same method on RNA-seq data from The Mount Sinai Brain Bank (MSBB) Study within the same research (https://www.synapse.org/Synapse:syn10526259). For validation of proteome data, we used the TMT-MS results from The Accelerating Medicines Partnership Alzheimer’s Disease Diverse Cohorts Study (AMP-AD_DiverseCohorts) (https://www.synapse.org/Synapse:syn55249985). For cell type proportion analysis, we used snRNA-seq data from ROSMAP study participants (https://www.synapse.org/Synapse:syn31512863).

We used ChatGPT and DeepL for the assistance of manuscript editing and coding.

The authors thank Nancy R. Gough (BioSerendipity, LLC) for editorial assistance and critical input.

## Author contributions

T.K. and S.K. conceived and supervised the project; T.K. and H.S. analyzed the data; writing group consisted of T.K., K.M., H.S., H.W., and S.K.

## Supplementary figures

**Figure S1.**
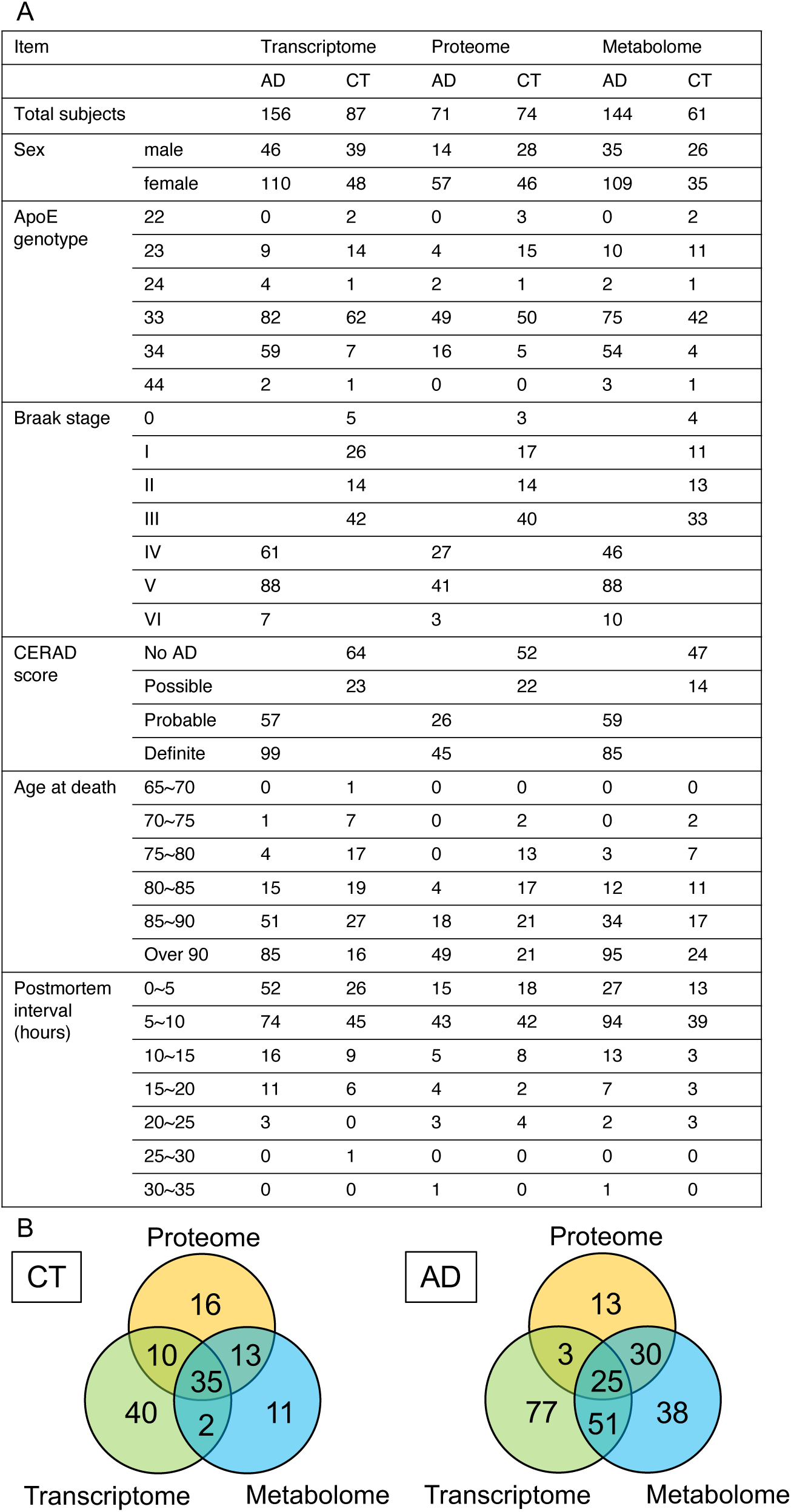
Information for the subjects from ROSMAP study. (A) Information of the samples from ROSMAP Study used in this study, including sample size, sex, ApoE genotype, Braak Stage, CERAD score, age at death, and postmortem interval (PMI). (B) Venn diagrams showing the overlap in data available for each subject across the three omic levels: transcriptome, proteome, and metabolome.

**Figure S2.**
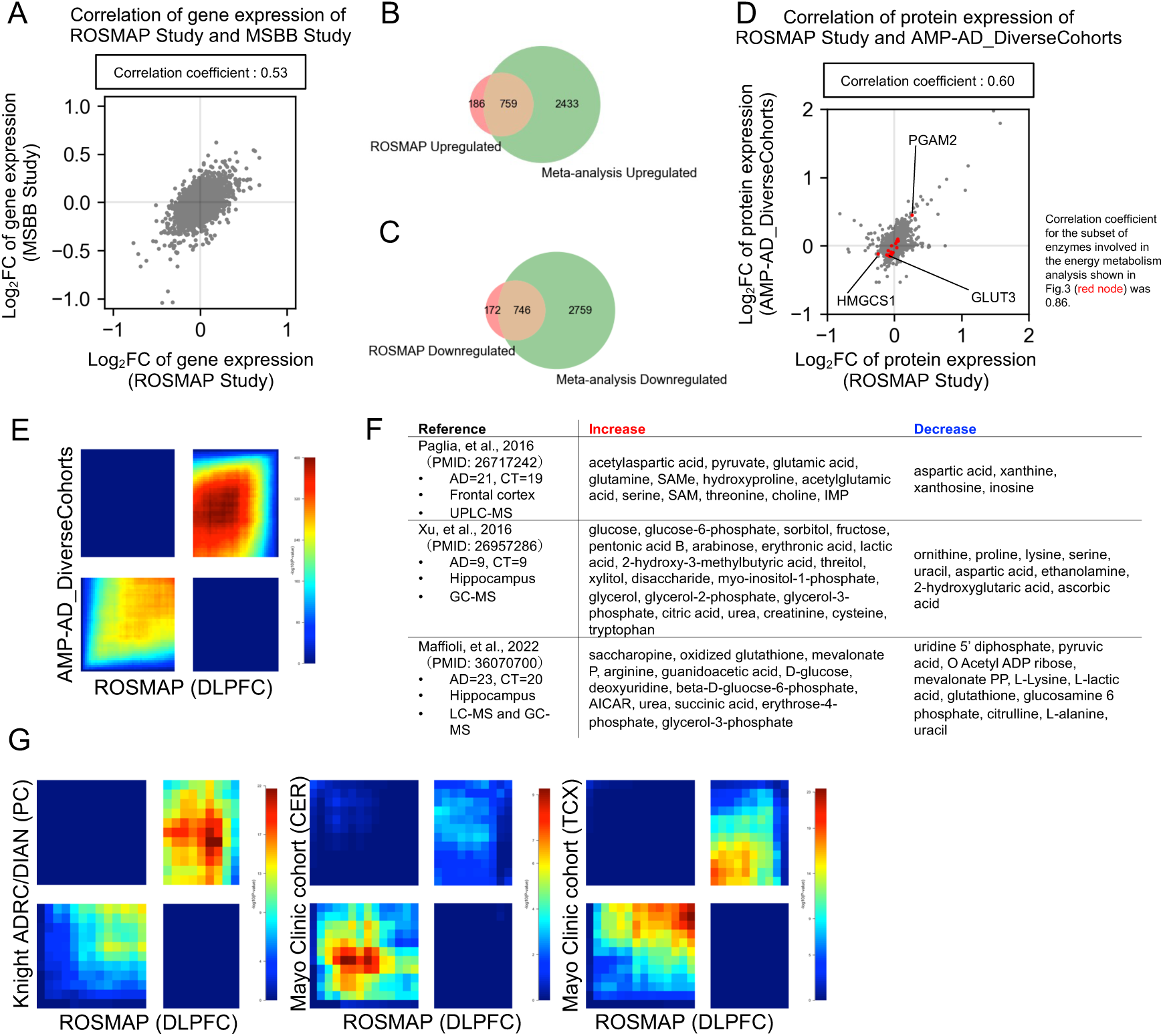
Comparison of transcriptome, proteome, and metabolome data from the ROSMAP Study with data from other studies. (A) Correlation between two transcriptomic datasets obtained from the ROSMAP Study and the MSSB Study for each gene. In the MSSB Study, RNA sequencing data were collected from brain tissue (FP; frontal pole) of individuals with AD (45 AD patients) and control subjects (45 CT subjects). For correlation analysis, the log_2_ fold-change values in 16,348 genes were calculated by dividing the average expression of the AD group by the average expression of the CT group. (B) Venn diagrams showing the overlap of upregulated DEGs between the ROSMAP study and those identified in a large-scale meta-analysis across seven brain regions [11]. (C) Venn diagrams showing the overlap of downregulated DEGs between the ROSMAP study and those identified in a large-scale meta-analysis across seven brain region [11]. (D) Correlation between two proteomic datasets obtained from the ROSMAP study and AMP-AD_DiverseCohorts study for each protein. TMT-MS data from each study were from brain tissue (DLPFC) of individuals with AD (637 AD patients) and control subjects (224 CT subjects). Correlation analysis was performed on the log_2_ fold-change values (average abundance of AD divided by average abundance of CT) for 9,180 proteins. Specifically, red nodes represent the subset of enzyme proteins which are involved in energy metabolism shown in Fig.3. (E) Rank-rank hypergeometric overlap (RRHO) heatmap comparing proteomic changes in AD compared to CT between the ROSMAP study (x-axis) and the AMP-AD_DiverseCohorts study (y-axis). Proteins were ranked based on -log_10_(q-value) multiplied by the sign of log_2_ fold-change in each dataset. Color intensity represents -log_10_(p-value) of the overlap at each rank threshold. Enrichment in bottom-left and top-right quadrants indicates concordant changes between the two datasets. (F) Differentially expressed metabolites (DEMs) identified in previous studies [1,2,3]. Note that each study used different definitions for AD and CT groups, had variations in measurement methods and brain regions analyzed. These studies had smaller sample sizes than our study. (G) RRHO heatmaps comparing metabolite changes in AD compared to CT between the ROSMAP study (DLPFC) (x-axis) and independent metabolomic datasets from multiple cohorts and brain regions [12,13] (y-axis). The Knight ADRC/DIAN dataset [13] consisted of 79 postmortem brain samples from cerebellar cortex (CER) of AD, 24 samples from CER of CT, 63 samples from temporal cortex (TCX) of AD, and 20 samples from TCX of CT and quantified 627 metabolites. The Mayo Clinic cohort datasets [12] consisted of 305 postmortem brain samples from parietal cortex (PC) of AD, 26 samples from PC of CT and quantified 685 metabolites. Metabolites were ranked based on -log_10_(p-value or q-value) multiplied by the sign of log_2_ fold-change in each dataset. Enrichment in bottom-left and top-right quadrants indicates concordant changes between the two datasets.

**Figure S3.**
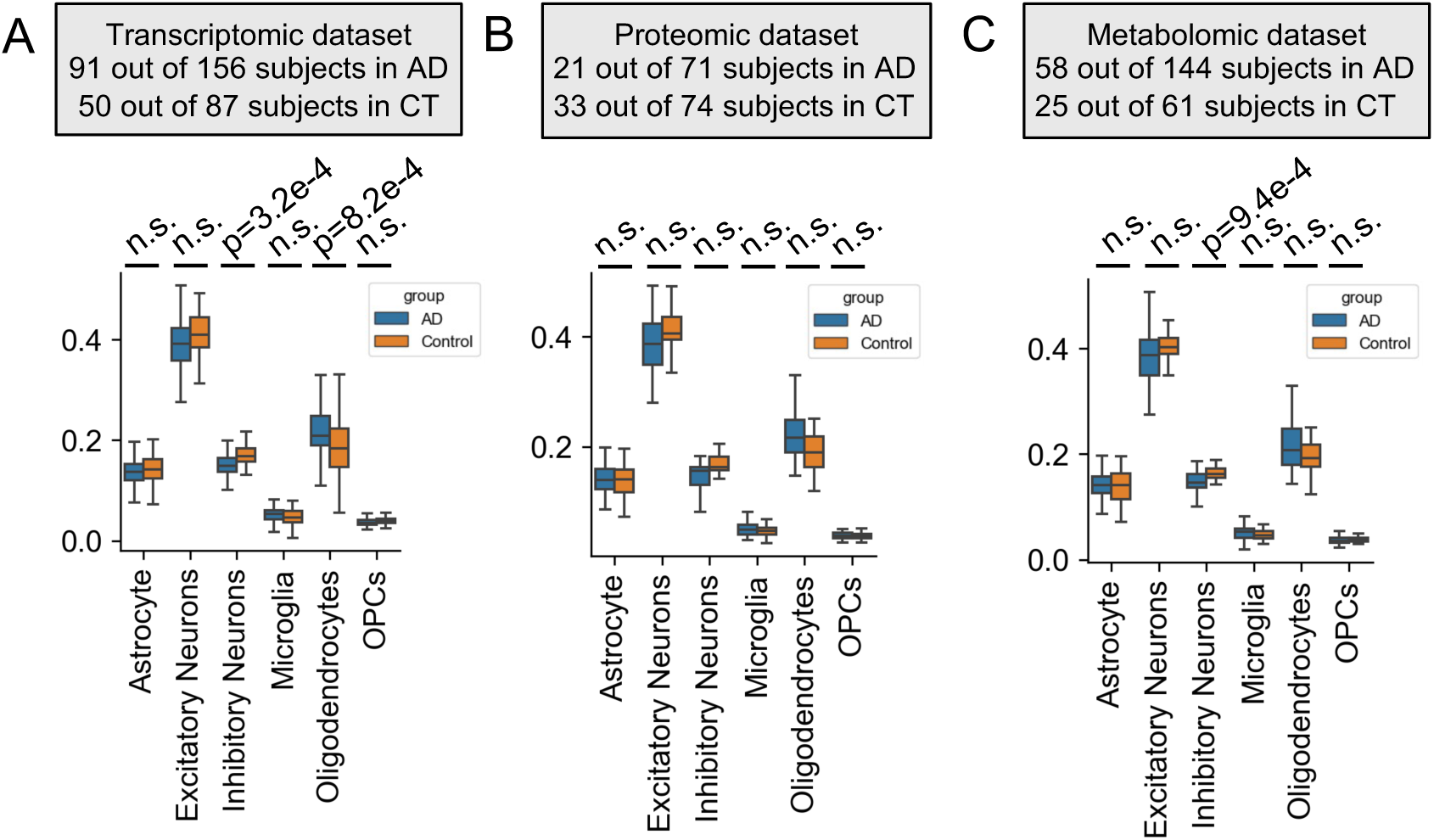
Cell type proportions estimation using a previous single-cell dataset. Cell type proportions were estimated using a publicly available single-nucleus transcriptomic dataset from DLPFC tissue of 465 subjects in the ROSMAP cohort [14]. For each omic dataset, overlapping subjects were identified and cell type proportions were calculated for the matched samples. Box plots show the proportions of astrocytes, excitatory neurons, inhibitory neurons, microglia, oligodendrocytes, and OPCs in the AD group (blue) and the control group (orange) for the transcriptomic dataset (A; n = 91 AD, n = 50 CT), the proteomic dataset (B; n = 21 AD, n = 33 CT), and the metabolomic dataset (C; n = 58 AD, n = 25 CT). Statistical comparisons were performed using two-tailed t-tests with Bonferroni correction (significance threshold: p-value < 8.3 × 10^-3^). n.s., not significant.

**Figure S4.**
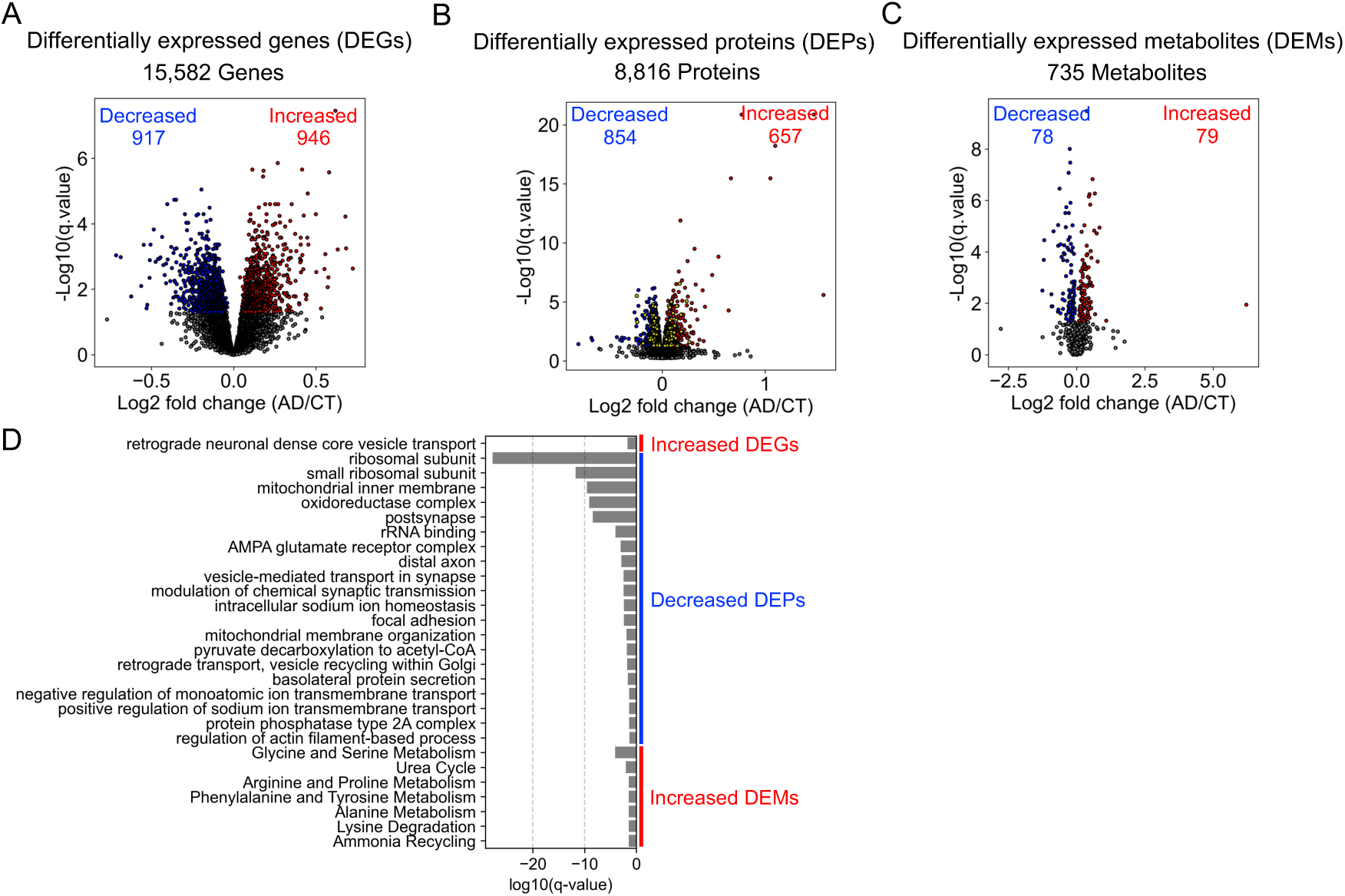
Differential and Enrichment analysis for differentially expressed molecules of AD. (A) Volcano plot for differentially expressed genes in AD. Increased mRNAs are shown in red. Decreased mRNAs are shown in blue. (B) Volcano plot for differentially expressed proteins in AD. Increased proteins are shown in red. Decreased proteins are shown in blue. Differentially expressed enzyme proteins are shown in yellow. (C) Volcano plot for differentially expressed metabolites in AD. Increased metabolites are shown in red. Decreased metabolites are shown in blue. (D) Enrichment analysis for DEGs, DEPs, and DEMs in AD performed with Metascape (for DEGs and DEPs) and MetaboAnalyst (for DEMs). Only significantly enriched terms (q-value < 0.05) are shown.

**Figure S5.**
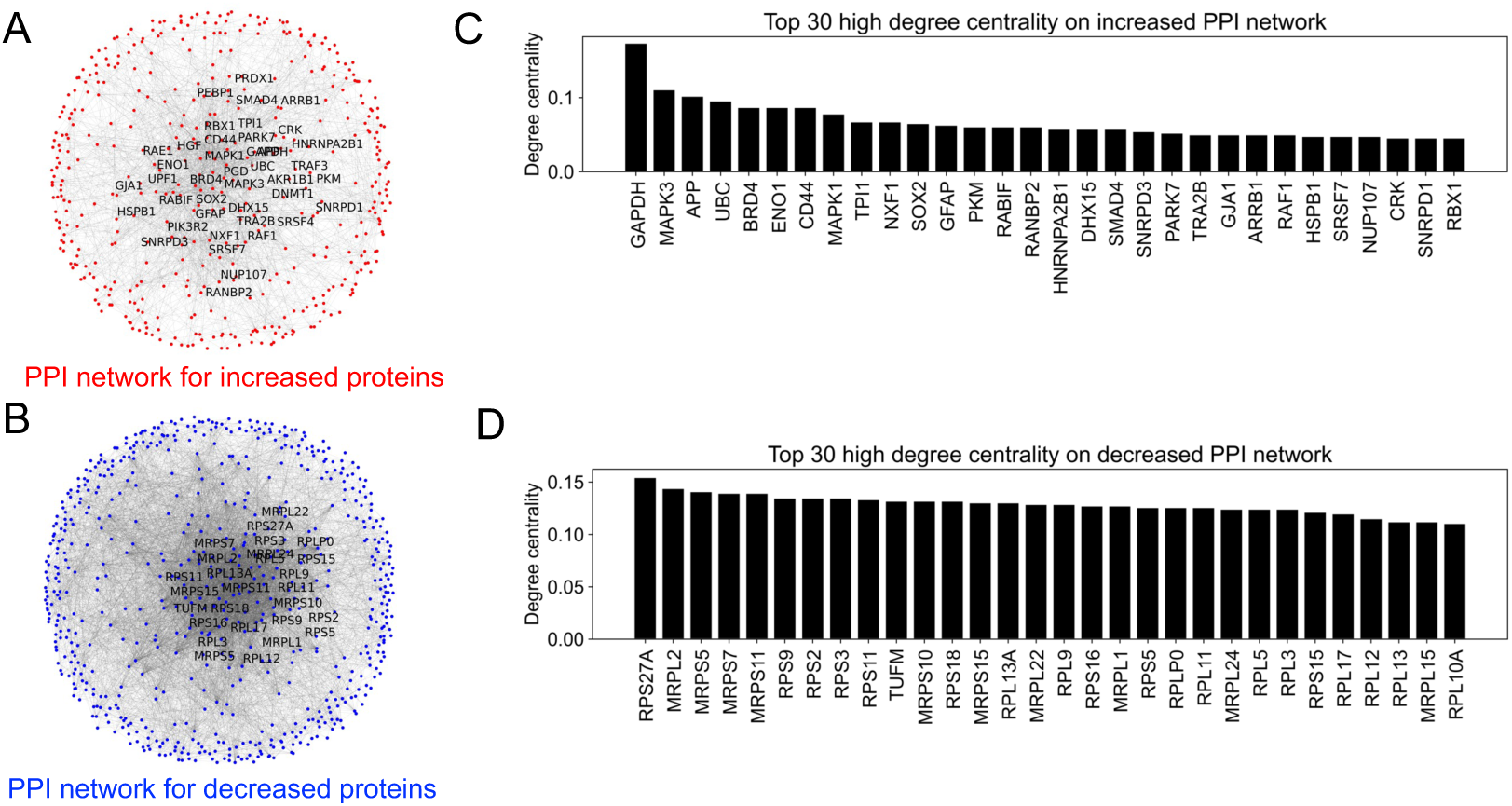
Construction of protein-protein interaction networks in DLPFC of AD. (A) The protein-protein interaction (PPI) network for increased DEPs. Red nodes and gray edges indicated DEPs and their interactions, respectively. The nodes whose degree are over 18 their degree centralities are labeled for reasons of legibility (Supplementary Data 2). (B) The PPI network for decreased DEPs. Blue nodes and gray edges indicated DEPs and their interactions, respectively. The nodes whose degree are over 75 their degree centralities are labeled for reasons of legibility (Supplementary Data 2). (C) Top 30 DEPs with high degree centrality on the increased PPI network. (D) Top 30 DEPs with high degree centrality on the decreased PPI network.

**Figure S6.**
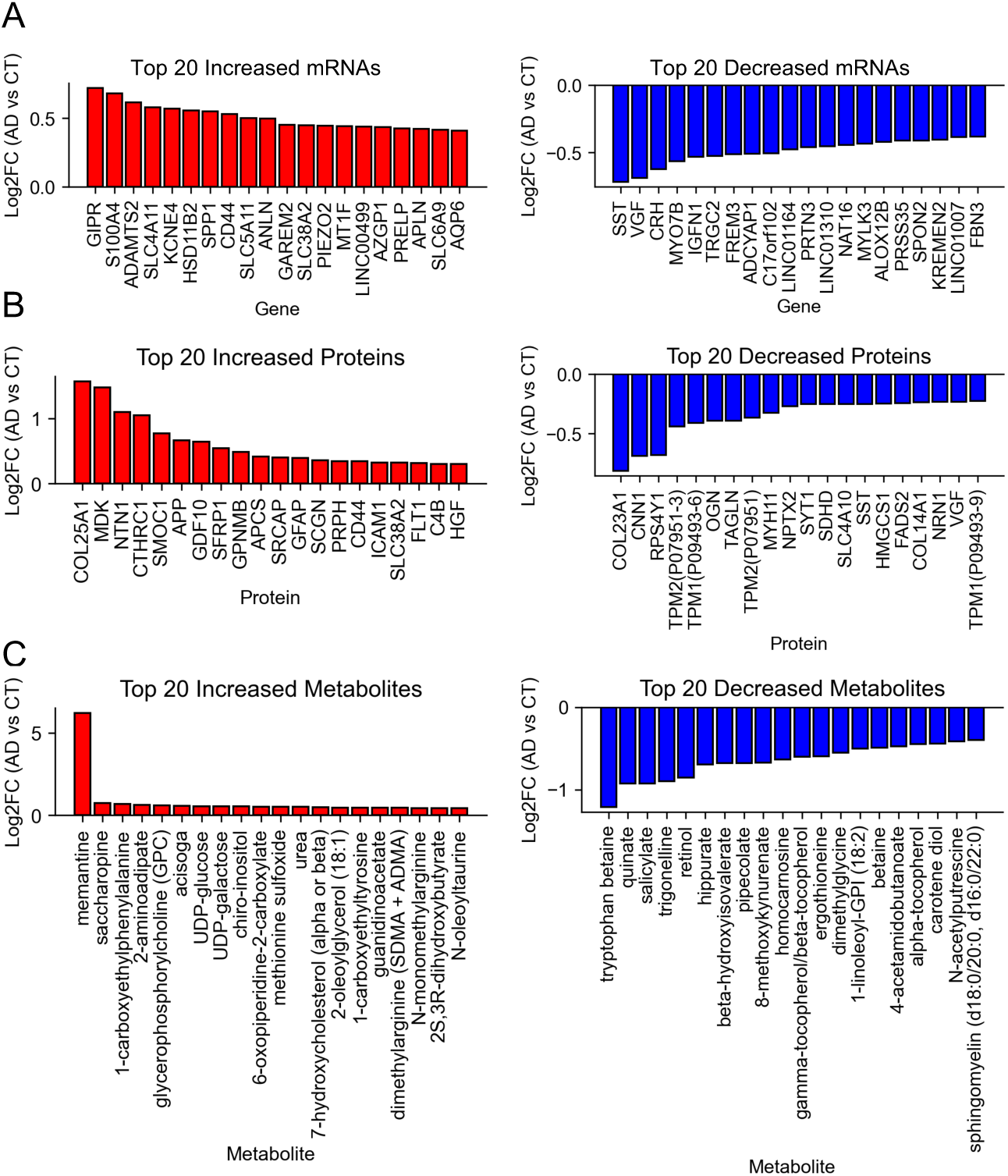
Top 20 differentially expressed genes, proteins, and metabolites. (A) The top 20 differentially expressed genes (DEGs) ranked and plotted according to log_2_ fold change (Log2FC) in mean expression between AD and CT. Red, DEGs with increased expression in AD (left); blue, DEGs with decreased expression in AD (right). (B) The top 20 differentially expressed proteins (DEPs) ranked and plotted according to log_2_ fold change (Log2FC) in mean expression between AD and CT. Red, DEPs with increased expression in AD (left); blue, DEPs with decreased expression in AD (right). (C) The top 20 differentially expressed metabolites (DEMs) ranked and plotted according to log_2_ fold change (Log2FC) in mean expression between AD and CT. Red, DEMs with increased expression in AD (left); blue, DEMs with decreased expression in AD (right).

**Figure S7.**
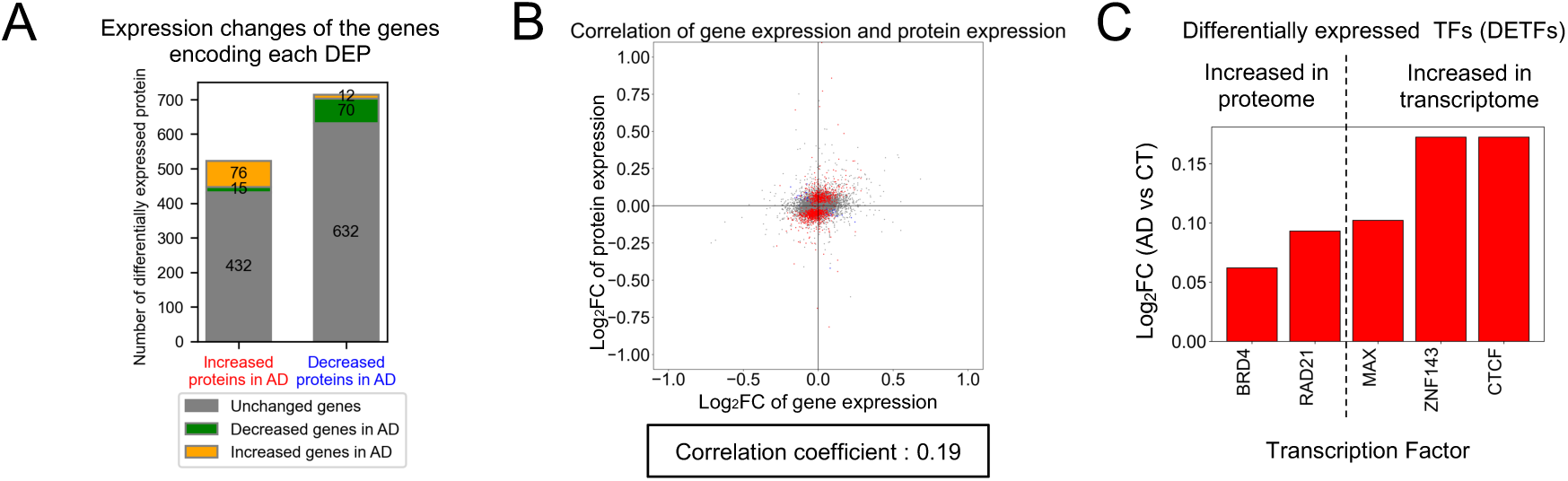
Identification of differential regulations connecting differentially expressed molecules across different omic layers. (A) The change in the encoding gene for the corresponding DEP grouped according to the increased DEPs and decreased DEPs in AD. (B) Correlation of the log_2_ fold changes (AD group vs. CT group) of measured molecules at the transcriptome level (x-axis) and proteome level (y-axis). Each point on a scatter plot represents a molecule measured as both mRNA and protein. DEPs are shown in blue or red. Blue represents those DEPs with encoding DEGs that change in the opposite direction; Red represents the rest of the DEPs. (C) Log_2_ fold change (AD group vs. CT group) of each differentially expressed transcription factor (DETF). The log2 fold change (Log2FC) of gene expression (transcriptome) or protein expression (proteome) of the indicated DETFs, which were identified from the 19 TFs that were inferred as potential regulators of DEGs with significantly altered expression in AD.

**Figure S8.**
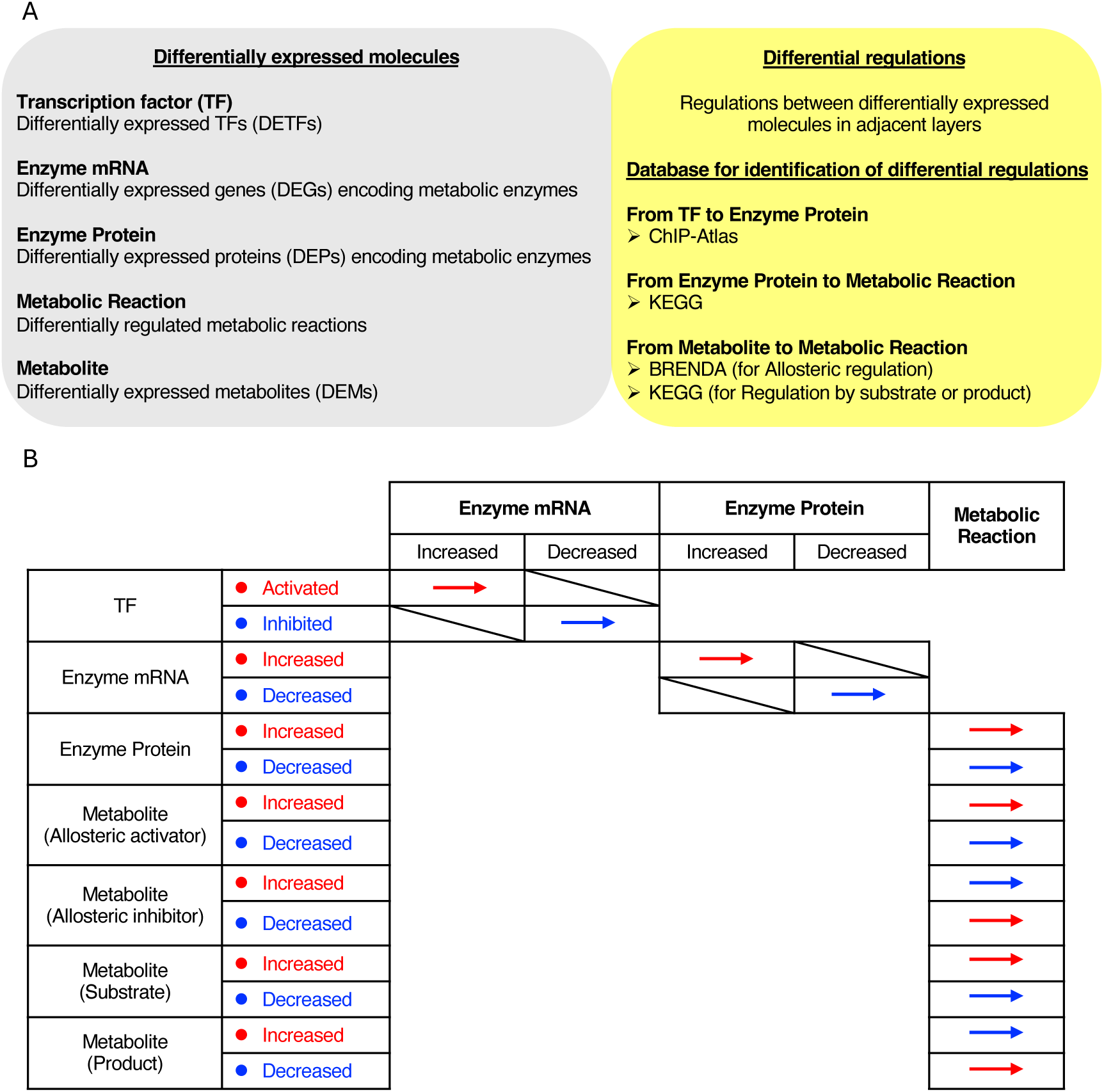
Contents of the trans-omic network for differentially regulated metabolic reactions between CT and AD. (A) The definition of differentially expressed molecules in each layer (left). Differential regulations between adjacent layers and the databases used to identify them (right). (B) Classification of the differential regulations (activating or inhibiting in AD group). The methodology of constructing differential regulatory trans-omic network of metabolic reactions is based on our previous study [4,5]. Briefly, differentially expressed TFs (DETFs), genes (enzyme mRNAs), proteins (metabolic enzymes), metabolic reactions, and metabolites were integrated to construct a five-layer network for AD. Adjacent layers were linked by known regulatory relationships. Edge colors indicate the direction and functional sign of changes: red/blue denote activating/inhibiting regulation for TF-mRNA, mRNA-protein, and protein-reaction edges. For reaction-metabolite links, edge colors were assigned based on the metabolite change and its role (allosteric activator/inhibitor or substrate/product).

## Supplementary text

### Top 20 differentially expressed genes, proteins, metabolites. (Fig. S6)

We identified the top 20 increased and top 20 decreased DEGs based on their log_2_ fold change (log2FC) values (AD group vs. CT group) for their mean values (Fig. S6A).

*GIPR*, which has been reported to have neuroprotective effects [6], and the inflammation-associated gene *CD44* were identified as DEGs with high log2FC values. *SST*, encoding the neuropeptide somatostatin, was identified among the DEGs with the lowest log2FC values.

We also identified the top 20 upregulated and top 20 downregulated DEPs based on their log_2_ fold change (log2FC) values (Fig. S6B). DEPs with high log2FC values included SIRPB1, which promotes phagocytosis by microglia [7], and GFAP, a candidate blood biomarker of AD [8]. DEPs with low log2FC values included those associated with the cytoskeleton, such as CNN1, TMP1, and TMP2.

We also identified the top 20 DEMs with the highest and lowest log2 fold change (log2FC) values (Fig. S6C). Notably, memantine was identified as the metabolite with the highest log2FC value. Memantine is a therapeutic drug for Alzheimer’s disease.

Therefore, it does not seem to be related to the pathology of the disease itself. However, this means that our analysis can accurately detect the differences between CT and AD groups. Retinol and sphingomyelin showed the lowest log2FC values.

### Protein-protein interaction networks for the DEPs in AD. (Fig. S5)

The multi-omic data used in this study contains not only metabolism, but also more general cellular mechanisms such as gene-regulatory network and protein-protein interactions. Therefore, we constructed protein-protein interaction (PPI) networks using DEPs with STRING [9] (Fig. S5A, S5B and Supplementary Data 2). We then examined the degree centralities of the nodes on the PPI networks (Fig. S5C and S5D). In the “Increased PPI” network, GAPDH emerged as the protein with the highest degree centrality. GAPDH is not only an enzyme in the glycolytic pathway but also plays roles in apoptosis, neurodegeneration, cognitive dysfunction, and the formation of aggregates with Aβ [10]. In contrast, in the “Decreased PPI” network, nearly all the top 30 proteins by degree centrality were ribosomal proteins or mitochondrial ribosomal proteins. These results suggested that AD is associated with reduced translation both in nucleus and mitochondria.

## Supplementary tables

**Table S1.** A list of transcription factors inferred to be involved in the regulation of DEGs in DLPFC of AD.

| Classification | TF Name |
| --- | --- |
| for Increased Gene | CTCF, MAX, MYCN, OLIG2, KDM1A, CTCFL, ARID1A, LIN28B, GTF3C2, CHD8, SMC3, ZNF92, TOP2B, ZNF143, EP300, TEAD4, BRD4, RAD21, BRD2 |
| for Decreased Gene | None |

**Table S2.** A list of differentially expressed metabolic enzymes in DLPFC of AD.

|  |  |
| --- | --- |
| Increased enzymes | ACAT2, AK1, AKR1B1, ALDH3A2, AMPD2, ASL, B4GALNT1, BBOX1, BLVRA, BLVRB, CA3, CAD, CBR3, CBS, CMBL, DGKG, DNMT1, ENO1, ESD, FAH, GALK1, GALM, GALNT16, GAPDH, GBE1, GMPR, GSTP1, INPP4B, ITPKB, LPIN1, LTA4H, MAN2A2, MAOB, MTAP, MTHFD1, MTHFD2L, NAGK, NT5C, NUDT16, OCRL, PAFAH1B3, PGAM2, PGD, PGLS, PHYKPL, PI4K2A, PKM, PLCD1, PLCD3, PPT1, PRDX6, QPRT, RBKS, RENBP, SAT2, SORD, TPI1, UGP2, UROD |
| Decreased enzymes | ABHD16A, ACAA2, ACOX1, ACSS1, ADCY1, ADCY2, AGPAT1, AGPAT3, AHCYL1, ALDH4A1, ALPL, AOC3, BCAT2, BCKDHB, BDH1, BPNT2, CDIPT, CS, CTH, DBT, DGKH, DGKI, DGKZ, DLD, ECHS1, FADS2, FDFT1, GAD1, GAD2, GGT5, GK, GLDC, GPHN, GSTK1, HMGCL, HMGCS1, HMOX2, HSD17B12, IDH2, L2HGDH, LCLAT1, MBOAT2, MECR, MGLL, MGST3, MT-ND3, MTMR7, NAT8L, NDUFS1, NDUFS7, NDUFS8, NDUFV1, NFS1, NNT, NOS1, NT5E, PDE4A, PDE4B, PGM2L1, PIP5K1A, PIP5K1C, PLCB1, PRPS2, PYCR2, RRM1, SDHA, SETD7, SHMT2, SMPD3, TM7SF2 |

## Supplementary data

Supplementary_Data_1.xlsx: Trans-Omic network for metabolic reactions in AD

Supplementary_Data_2.xlsx: Protein-protein interaction networks in AD

